# A portable and high-integrated 3D microfluidic chip for bacterial quantification and antibiotic susceptibility testing

**DOI:** 10.1101/2021.11.22.469235

**Authors:** Wenshuai Wu, Gaozhe Cai, Yang Liu

## Abstract

On-site single-cell antibiotic susceptibility testing (sc-AST) provides unprecedented technical potential to improve the treatment of bacterial infections and study heterogeneous resistance to antibiotics. Herein, we developed a portable and high-integrated 3D polydimethylsiloxane (PDMS) chip to perform fast and on-site bacteria quantification and sc-AST. The 3D arrangement of the chambers significantly improved the integration of reaction units (∼10000/cm^2^) and widened the dynamic range to 5 orders of magnitude. A capillary valve-based flow distributor was adopted for flow equidistribution in 64 parallel channels and uniform sample loading in as short as 2 s. The degassed PDMS enabled this device to independently dispense the sample into 3D chamber array with almost 100% efficiency. The quantification of *Escherichia coli* (*E. coli*) strains with various activity was accomplished in 0.5-2 h, shortened by 20 h in comparison to the traditional plate counting. The functionality of our platform was demonstrated with several effective antibiotics by determining minimum inhibitory concentrations at single-cell level. Furthermore, we utilized the lyophilization of test reagents and needle-mediated reagents rehydration to realize one-step on-site sc-AST. The results indicate that the proposed sc-AST platform is portable, highly sensitive, fast, accurate and user-friendly, thus it has the potential to facilitate precise therapy in time and monitor the treatment. Meanwhile, it could serve as an approach for investigating the mechanisms of heteroresistance at single-cell resolution.

## INTRODUCTION

Bacterial infections have become a significant threaten to human health worldwide and the emergence of antibiotic resistance aggregates the difficulty and cost for fighting against bacterial infections.^1^ The empirical therapies for infectious diseases with broad-spectrum antibiotics or excessive dosage make the situation progressively worse and further lead to the generation of multidrug-resistant pathogens.^2^ To avoid the improper treatment of bacterial infectious diseases, antibiotic susceptibility testing (AST) is conducted to help to select effective drugs and determine the minimum inhibitory concentrations (MIC) for individual patient. ^3^

Conventional AST, which contains broth dilution and disk diffusion, relies on massively cell growth to form visible turbidity or colonies and usually takes 16-20 h to report the susceptibility profiles of the pathogens.^4, 5^ The high amount of the cells required in these methods also leads to long preculturing time (taking 5–6 h) before AST.^5^ The long period of AST is incompatible with desired timely treatment of bacterial infections.^6^ In addition, the operation of standard AST depends on centralized clinical microbiology laboratory and specialized persons, hampering its application in resource-limited settings.^7^ In response, many microfluidic-based methods were developed to accelerate the process of AST. Among them, the strategies to indicate the cell resistance to antibiotics include the identification of resistant genes, ^8-10^ the cell growth monitoring^11-15^ and the metabolic activity sensing.^16-21^

The detection of metabolic activity of cells exposed to antibiotics is widely applied to perform fast AST in microfluidic devices. The bacteria are loaded in micro-sized chamber array^16-19^ or water plugs^22, 23^ in capillary tube. The fluorescence probe, resazurin, is employed for sensing the activity of cell respiration in the presence of antibiotics. The time required for resazurin-based AST has been pushed to less than 5h, which depends on the initial cell number in microreactors. To make the resazurin-based AST suitable for on-site analysis, researchers have developed several portable PDMS devices to conduct AST, in which the sample distribution was realized by pipette,^24^ syringe^21^ or vacuum stored in PDMS.^25^ These experiments have illustrated the potential of resazurin-based assay in fast and on-site AST. However, the proposed resazurin-based assays are not sensitive enough to detect bacteria that occur at low concentrations (less than 10^5^ CFU/mL) in clinical samples. The preculturing step to multiply the cell numbers bottlenecks the complete AST process and makes the existing methods fail to evaluate the infections burden.

The increasing studies have proved the existence of heterogeneous resistance that the bacteria in a population have different degrees of resistance to antibiotics.^26-29^ However, in the standardized AST, because of the low sensitivity and low throughput, the bacteria in original samples are firstly cultivated on agar and then several colonies are tested to represent the susceptibility profiles of the bacteria population.^5, 26^ Given the heterogeneous resistance, there is a consensus that the bacterial population in original samples should be analyzed directly for comprehensive AST in sync with the quantification of the bacteria load to estimate drug resistance and efficacy. ^5, 7, 26, 30^ Long turnaround time, low sensitivity and the lack of bacterial quantification drive researchers to develop innovative methods to meet the urgent and strict requirements.

To realize directly comprehensive AST and quantification on bacterial population in original samples, as well as on-site monitor the infections burden, new methodologies should have the characteristics of high sensitivity, broad dynamic range, short detection time, accurate quantification and portability. Fortunately, the single cell AST (sc-AST) based on microfluidic droplet technology offers the hope of comprehensive AST alongside the bacteria quantification.^28, 31-33^ With the scalability of droplet reactors, droplet-based sc-AST have broad dynamic range to handle the various clinical samples. Moreover, the droplet-based sc-AST is able to characterize the heteroresistance at single cell resolution and has potential in elucidating the mechanisms of heteroresistance.^27, 28^ However, the droplets generation relies on bulk pumps and the complex manual operation, which hinder its application for on-site analysis.^33,34^ Besides, the coalescence of droplets during incubation will impair the accuracy of sc-AST. The chamber-based microfluidic chip could be an alternative to solve these problems, but suffers from low chamber integration, leading to low throughput and narrow dynamic range.

In this study, we presented a portable and high-integrated chamber-based PDMS chip for fast and on-site bacteria quantification and sc-AST. Unlike the reported devices with 2D chamber array,^35, 36^ the 3D arrangement of the chambers was developed to increase the chamber density to ∼10000/cm^2^. The highly integrated chip has a wide dynamic range of 5 orders of magnitude. The capillary valve-based flow distributor was applied to scale up the channels from 1 to 64 and realize flow equidistribution. This developed PDMS chip has the ability of automatic sample loading and compartmentalization without the requirement of external instruments. The sample dispersion can be accomplished in 2 s with a customized needle, much faster than droplet generation. Using this device, we developed a digital resazurin assay to determine the concentration of different *E. coli* strains in 0.5-2 h and rapid sc-AST (< 4 h) for the selection of effective antibiotics and the detection of MICs. Besides, we investigated the pre-storage of test reagents in chambers by lyophilization and developed loss-free rehydration of dried reagents by needle-mediated sample loading, making it suitable for on-site sc-AST.

## RESULTS AND DISCUSSIONS

### Chip design

To improve the integration of chambers, reducing the interval between chambers is a common strategy but it will reach to the limit when the distance drops to 20 μm (Figure S1). Here, we firstly extended the chamber arrangement to 3D scale. The design of 3D chamber array breaks through the limitations without increasing the complexity in fabrication. As shown in Figure 1a, the microfluidic chip with the size of 5 cm by 1.8 cm consists of two PDMS layers and a glass slide. Each PDMS layer has channel network for sample distribution and 2D chamber array (Figure 1b). The height of channel and chambers are 20 μm and 45 μm, respectively. After assembled, the chambers in the upper layer are attached to the blank area between the chambers in the lower layer, forming the 3D chamber array with the interval of 20 μm (Figure 1c i and ii). In assembled chip, the main channels on these two layers are well aligned and form a closed channels with the depth of 40 μm (Figure 1c iii). The developed chip has 22660 chambers (50 μm) with the density of ∼10000/cm^2^, giving a broad dynamic range from 3.9 × 10^2^ to 8.9 × 10^7^ CFU/mL. With PDMS aligners reported in previous work,^37, 38^ the interval could be as small as 10 μm to further increase the chamber density. To introduce the sample solution into 3D chamber array, a bidirectional dispensing network was adopted, in which the inlet is located in the middle of the chamber array and the liquid flows in two directions simultaneously. Compared to the unidirectional distribution network,^35, 39^ the bidirectional one has a half of the path distance of liquid flow in the former. The shortened channels reduce flow resistance and save the time for sample loading. We developed a new flow distributor based on capillary valve to achieve fast and uniform sample distribution from 1 inlet to 64 parallel channels, which is a part of bidirectional dispensing network (Figure 1c i). The mechanism and performance of the dispenser will be discussed in next section.

**Figure 1.**
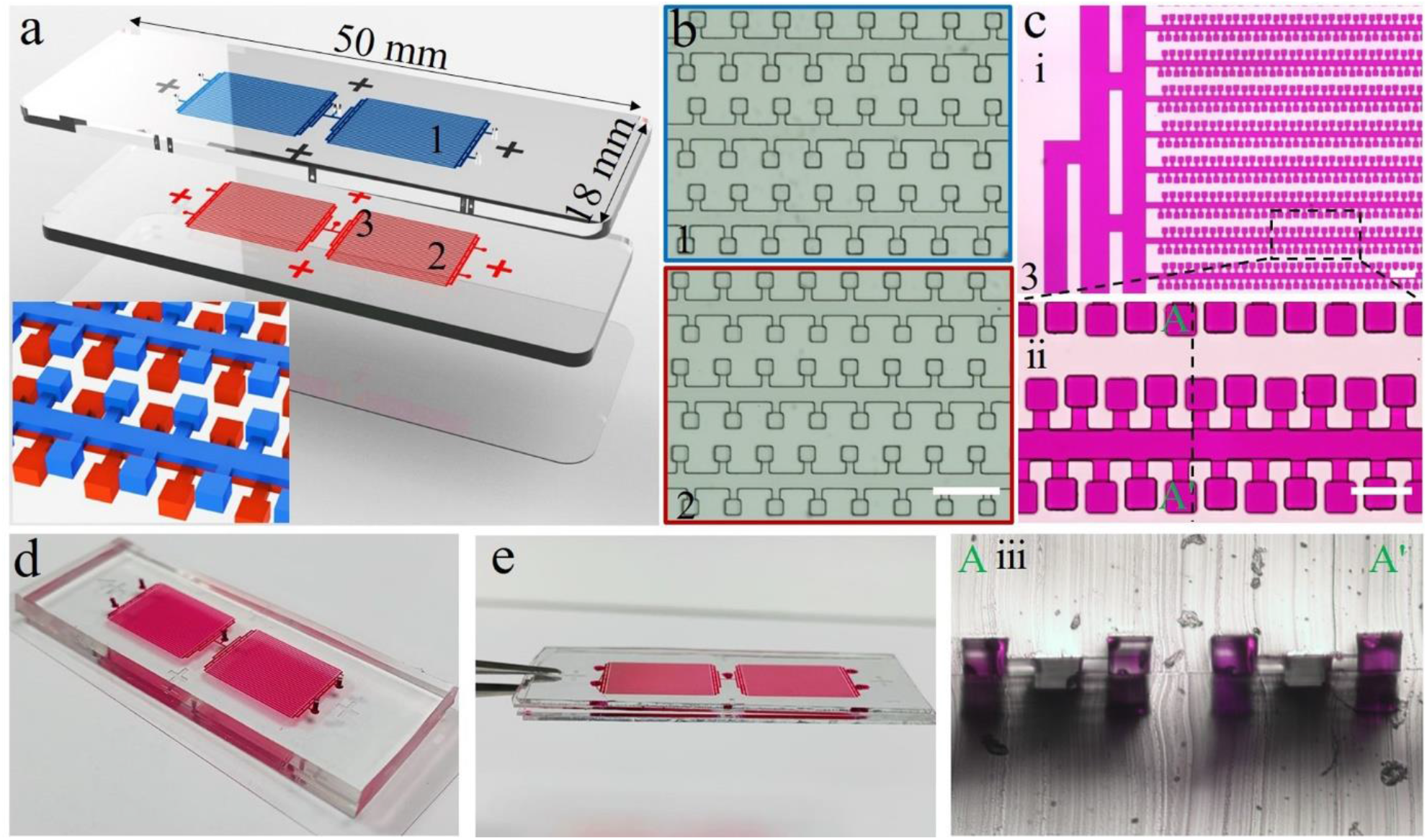
The high-integrated 3D chip for bacteria quantification and sc-AST. a) Exploded view of the 3D chip. The 3D chip consists of 2 PDMS layers and a glass layer. The 2 PDMS layers is bonded together face-to-face to form the 3D chip. The insert shows the 3D structures of the chamber array and channels after chip bonding. b) The channel and chamber geometries in upper (1) and lower layer (2). Scale bar denotes 200 μm. c) Dye-loaded chip for visualization of flow distributor and 3D chamber array (i). ii) an enlarged area showing 3D chamber array. iii) cross-section of 3D chamber array. Scale bar denotes 200 (i) and 100 μm (ii). d) a fabricated chip with 5.5 mm thickness. e) An ultra-thin chip (1.8 mm) for preventing from water evaporation. The thickness of the PDMS layer is 0.8 mm.

In this work, we constructed two types of chips, a normal chip and an ultra-thin chip (Figure 1d and e). Although the air permeability of PDMS contributes to independent and automatic sample distribution, it will cause water evaporation during incubation. The normal chip with 5.5 mm PDMS consumes more water to reach saturated vapor pressure, leading to water evaporation of peripheral chambers (Figure S2a). To solve this problem, an ultra-thin chip with 0.8 mm PDMS was fabricated. The glass slips (0.5 mm) on the top and bottom as permeation barriers confine the steam in the thin PDMS layer, ensuring saturated vapor pressure after slight evaporation (Figure S2b). Hydration channels surrounding the chamber array was reported to successfully reduce water evaporation of peripheral chambers.^36, 40^ The combination of hydration channels and ultra-thin PDMS may eliminate the water evaporation of all chambers. We also observed that putting the chip in DI water or in humid environment during incubation can mitigate the water loss. In the following experiment, the normal chip was used and incubated in humid environment.

### Capillary valve-based flow distributor

In reported studies, bifurcation method was successful in flow equidistribution and frequently used for scale-up of multiple channels. However, the flow impacts and turning losses in bifurcation system reduce the flow rate as the increasement of junction number. In addition, the bifurcation method increases the path distance of liquid flowing. The features limit the scale-up of channels. Therefore, a capillary valve-based flow distributor was developed and deployed in the negative pressure-driven chip to realize uniform sample loading in parallel channels. Figure 2a is a schematic illustration of the mechanical analysis of the distributor, mainly involving the parameters as the pressure and the surface tension. The Hydrodynamic resistance and the surface tension caused pressure drop, respectively denoted as *P*_*h*_ and *P*_*c*_. They contributed to the total pressure drop *P*_*t*_, i.e. *P*_*t*_ = *P*_*h*_ + *P*_*c*_.

**Figure 2.**
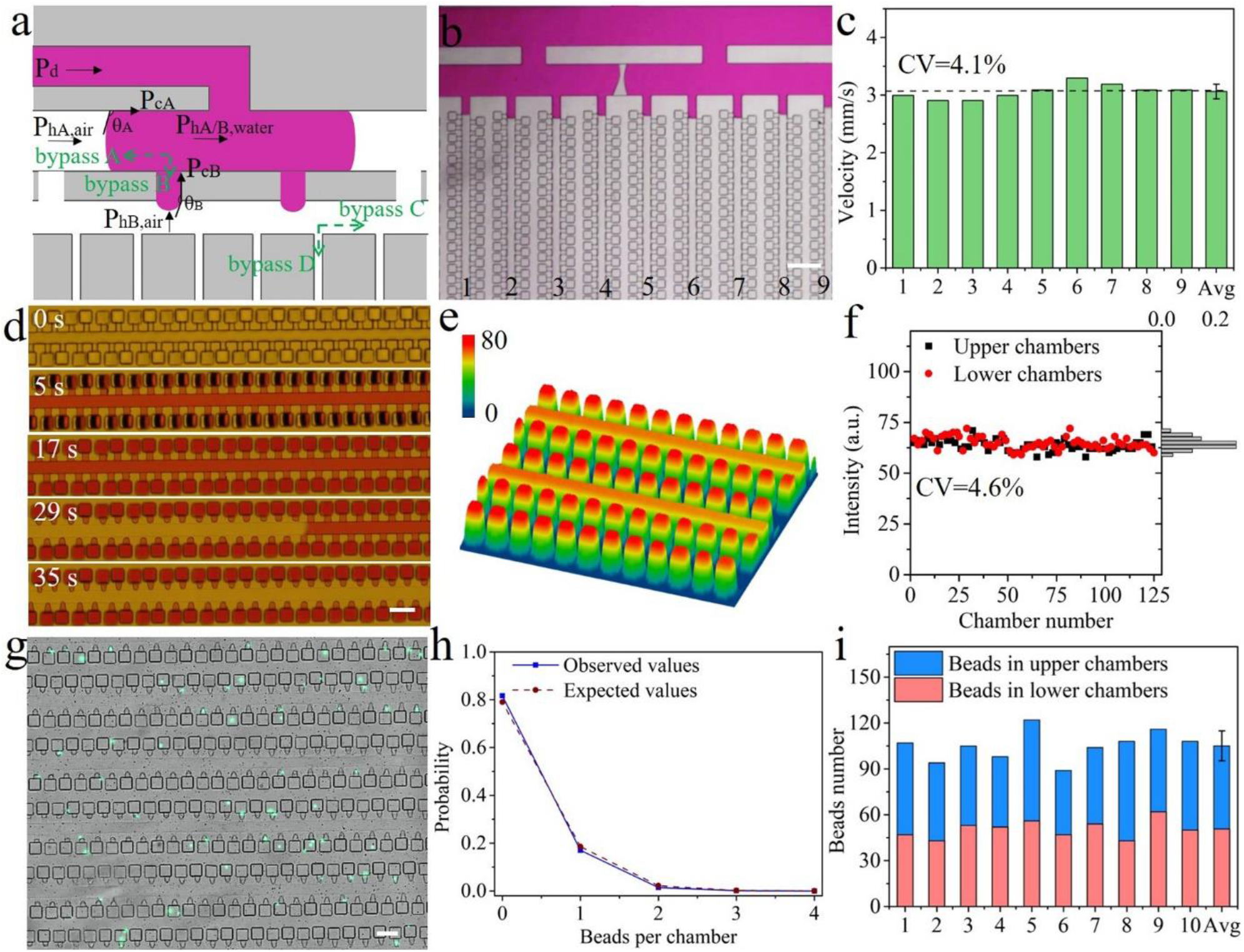
Characterizations of the high-integrated 3D chip. a) The mechanical analysis of liquid flow in the capillary valve-based flow distributor. b) The quasi-parallel filling of the vertical channels. Scale bar denotes 200 μm. c) The flow equidistribution in the parallel channels. The analyzed channels are that in (b). d) The process of sample distribution in 3D chamber array. Scale bar denotes 100 μm. e) The 3D intensity plot of chamber array with resorufin. f) The uniform fluorescence intensity of the chambers in different PDMS layers. g) The fluorescence photo of beads in chamber array. The bead concentration was 1.8 × 10^6^/mL, corresponding to 0.2 beads per chamber. h) The probability of cell number per chamber. i) Spatial distribution of beads in 10 random sections with 500 chambers. Error bar represents on the standard deviation of bead numbers in 10 sections.

Firstly, *P*_*h*_ can be calculated by the following equation,

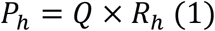

where, Q is the flow rate and *R*_*h*_ is the hydrodynamic resistance. For a Hele-Shaw microfluidic channel as used in our study, *R*_*h*_ can be calculated in the following way,

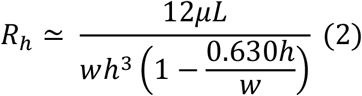

where, *μ* is the viscosity, *L* is the length of the channel, *w* is the width of the channel and *h* is the height of the channel.

The second term *P*_*c*_ has the origin of capillary force, which is calculated by the Young-Laplace equation in a rectangular situation,

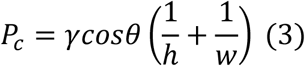

where, *γ* is the surface tension and *θ* is the contact angle.

Given the fact that the surface is hydrophobic (contact angle with water: 107 °), the capillary force impedes the sample flow. The driving force of the sample loading is the negative pressure in degassed PDMS. The flowing liquid encountered two bypasses, labeled A and B for each conjunction point with respectively high and low channel widths. Since *w*_*A*_ > *w*_*B*_ and *h*_*A*_ = *h*_*B*_, |*P*_*hB,water*_| > |*P*_*hA,water*_| and |*P*_*c,B*_| > |*P*_*c,A*_|, i.e., the resistance that sample liquid encounters in the bypass B is higher than the bypass A. Therefore, the mass flow rate of the bypass B is much smaller than the bypass A. After the bypass B is filled, the surface tension increases further more as the growing air-water interface at the end of bypass B will lead to an increasing contact angle (*θ*_b_) in a short period before contacting with the other side of the channel.

The previous analysis also applies to bypass C and D, and predicts a quasi-parallel filling of the vertical channels as the liquid prefers to fill the entire horizontal channel and then flows along the vertical one synchronically. It is supposed that the further scale-up of vertical channels would not obviously increase flow impacts, turning losses and the path length of liquid flow, which slow the liquid down, showing the superior performance to the bifurcation method. In the experiment, we indeed observed the quasi-parallel filling of the vertical channels after horizontal one was filled (Figure 2b and Video S1). In our analysis, the pressure drop due to air (*P*_*hA,air*_ and *P*_*hB,air*_) is not considered as the viscosity of air (1.813 × 10^−2^ mPa·s) is negligible comparing to that of water. The resistance of sample liquid in the vertical channels are almost the same once they are filled, thus theoretically the liquid moves in a similar speed with the same driving pressure. We extracted the velocity of the liquid in 9 channels based on the analysis of Video S1. The average velocity of them was 3.06 mm/s with a small coefficient of variation (CV = 4.1%), demonstrating the flow equidistribution in the parallel channels (Figure 2c). The mechanical analysis and the experimental results indicate the practicability of the developed flow distributor and a promising application in massive scale-up of the parallel channels.

### Self-driven sample loading

The chip operation and the principle of self-driven sample loading are shown in Figure S3. The negative pressure in degassed PDMS serves as the internal power to drive the liquid distribution and oil isolation (Figure 2d, Video S2). Two pieces of PDMS were attached to outlets to speed up the process of sample loading. The sample was loaded by a tip and subsequently sucked into chambers automatically till all of them were filled. Then, a thermosetting oil was introduced to get rid of aqueous from channels and partition the chambers. The time of tip-mediated sample dispersion depends on the volume of the liquid. It took 2 min, 3 min and 11 min for 8 μL, 10 μL and 20 μL, respectively. More importantly, the efficiency of sample distribution could reach to almost 100% under the optimized condition of sample volume and degassing time because of the uniform negative pressure in the 3D array. With vacuum packing,^41, 42^ the negative pressure in chip would be maintained for several month and execute self-powered sample partition without the requirement of external instruments, making the chip suitable for on-site analysis.

### The fluorescence detection of 3D chamber array

We firstly measured the size of chambers and assessed the uniformity of chamber volume, which are the important consideration for accurate determination of *E. coli*. The measured height and area of chambers are 45.2 μm (CV = 1.76%) and 2469 µm^2^ (CV = 1.76%), respectively (Figure S4). Therefore, the volume of chamber is 112 pL, which is calculated by multiplying the height and area of chamber. The small number of CV for chamber height and area indicates the good uniformity of the chamber volume. The 3D chamber array with the height span of 90.4 μm may cause blurred and uneven fluorescence imaging of the chip. To validate the uniformity of the fluorescence imaging, the 3D chamber array was filled with resorufin solution and imaged using a 10× magnification objective lens (numerical aperture 0.3). The 3D surface plot in Figure 2e shows uniform intensity of all chambers. The fluorescence intensity of chambers was in a narrow range with a CV value of 4.6% (Figure 2f). Given the small variability of the chamber volumes, the difference of the fluorescence intensity caused by imaging could be smaller. When comparing the average intensity of upper array with that of the lower one in one picture, there was no significant difference, indicating the uniform fluorescence imaging of 3D chamber array can be achieved (Figure S5). Therefore, in this work, 10× lens with 0.3 N.A. was applied to observe and capture the fluorescein in chambers.

### The random distribution of *E. coli* in the 3D chamber array

A precondition of digital quantification of *E. coli* is the stochastic distribution of single targets in chambers and then Poisson distribution can be used for calculating the accurate cell concentration. We used polystyrene (PS) beads with green emission to simulate the distribution of *E. coli* in the 3D chamber array. PS beads possess similar size (2 μm) and density (1.05 g/cm^3^) to *E. coli*. As indicated in Figure 2g, when the average numbers of cells per chamber (λ) was 0.2, most chambers were empty and ∼16.3% chambers had 1 bead. There were ∼1.9% chambers with more than 1 beads. The observed probability for beads number per chamber was in agreement with Poisson distribution (red line), indicating that *E. coli* would be discretized in chamber array randomly (Figure 2h). Moreover, the spatial amounts of beads in 10 random sections (500 chambers per section) were counted. The beads were classified into two groups according to the layers that they were in. In the case of 0.2 beads per chamber, in principle, there should be 100 beads in a section and half of them in one side of the 3D array. As shown in Figure 2i, a section had 105 beads on average, close to the theoretical value. The small error bar revalidated the random and uniform distribution of beads. Surprisingly, the beads tended to go into upper chambers, leading to more beads in upper layer (∼55 beads per section on average). Although there was the tendency of bead distribution between upper and lower layer, the distribution of bead in the 3D chamber array still conformed to Poisson distribution. Therefore, it has less effect on the quantification of *E. coli*. Despite that, we further confirmed that the design of 3D chamber array has less influence on bead distribution in principle by simulation (Figure S6). The difference of the bead capture between upper and lower chambers may result from the thicker PDMS of upper layer, which could make the chambers in it keep more solution and thus increase the amount of bead.

### Time-lapse imaging of *E. coil* detection course

To realize fast AST in sync accurate quantification of *E. coli*, a digital resazurin assay was developed in 3D array chip (Figure 3a). Resazurin, a cell permeable redox chemical, can be reduced into the fluorescent resorufin (560/590 nm) by viable *E. coli* to indicate the metabolic activity of cells. The signal of resorufin generates with cells growth and is proportional to the number of metabolically active cells. By partitioning single cells in 112 pL chambers, the resorufin produced is confined and concentrated in such small volume and could reach to the detectable concentration in short period. The detection time is dependent on the assay conditions including resazurin concentration, temperature, and the type and proportion of medium. The reduction rate of resazurin attained its maximum at resazurin of 0.04 mM, 50% LB broth and the temperature of 42 °C (Figure S7). Given the limited volume of chambers, 0.07 mM resazurin and 100% LB were applied to supply the adequate substrate and nutrition.

**Figure 3.**
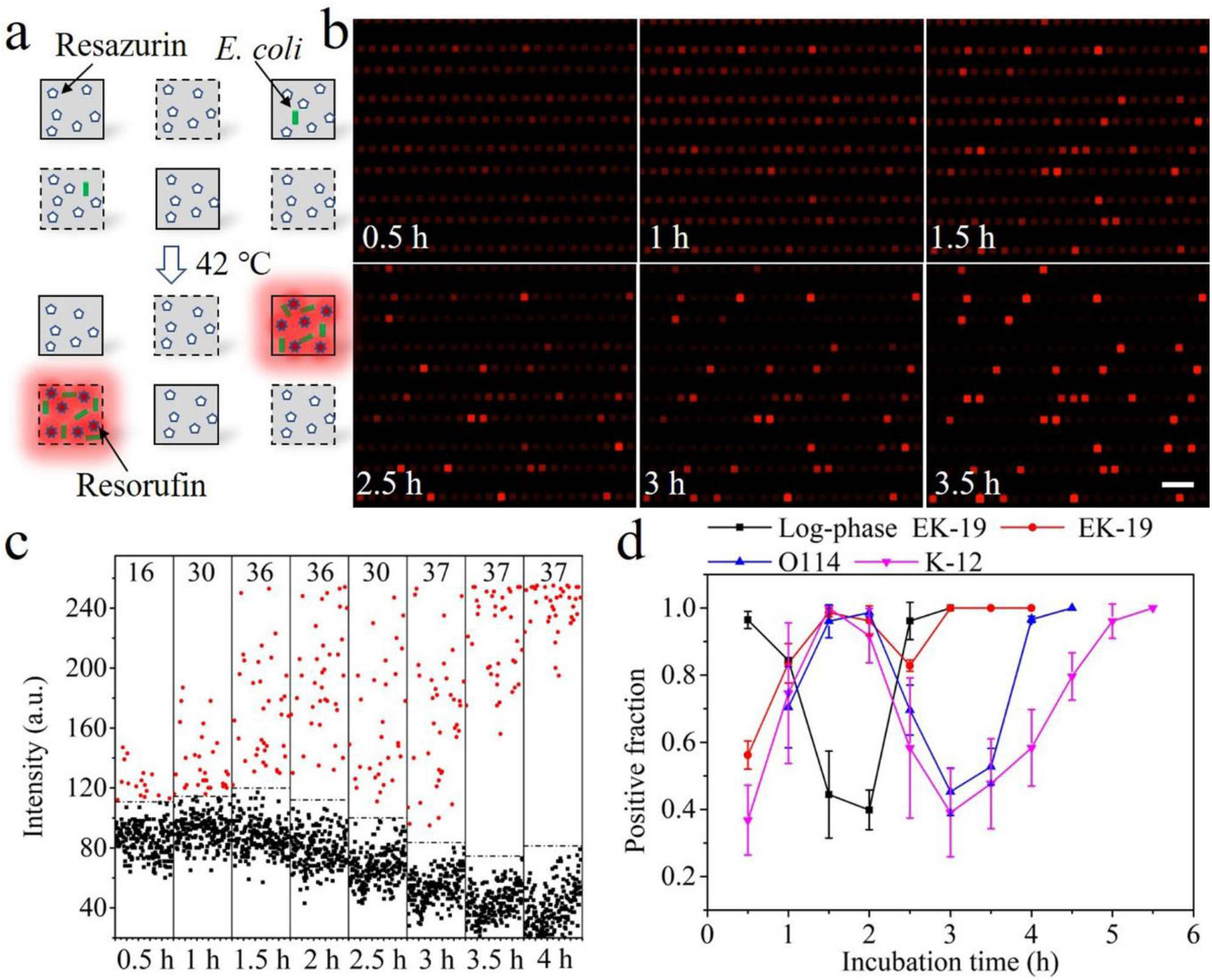
Time-lapse imaging of *E. coli* detection course. a) The concept of digital resazurin assay. b) The time-lapse imaging of stationary *E. coli* EK-19 detection. Scale bar denotes 200 μm. c) Fluorescence intensity of the chambers in (b) over time. The threshold equals the sum of average intensity of the chambers without cells and 3 times standard deviation. d) The evolution of positive fraction of multiple *E. coli* strains during incubation. The number of positive chambers at a certain time divided by the total number of positive chambers at the end of incubation was defined as the positive fraction.

In order to figure out the minimal detection cycle of single *E. coli* in chip, time-lapse imaging of *E. coil* detection was conducted by using Zeiss microscope equipped with an incubator. In this work, we tested an environmental strain, *E. coli* EK-19 at both stationery and log phase, and two lab strains, *E. coli* O114 and K-12. The sample solution containing stationary EK-19 (1 × 10^6^ CFU/mL, plate counting) were used, corresponding to 0.13 cells per chamber (λ). In this case, 93.4% positive chambers contained one *E. coli*. Representatively real-time images are shown in Figure 3b with a period of 0.5–3.5 h. The intensity of chambers was extracted and plotted in Figure 3c, visually reflecting the intensity of the compartments during incubation. The red dots above the threshold (dash line) were regarded as cell-containing chambers (positive). Roughly, the intensity of positive chambers became stronger over time. The number of positive chambers increased in 2 h and almost all the occupied chambers (98.6%) showed distinguishable fluorescence after 1.5 h incubation. Afterward, the fluctuation of the number of positive chambers occurred. All the chambers containing cells had sufficient fluorescence to be recognized at 3 h (Figure 3c and 3d). When the log phase EK-19 (OD 0.15) was loaded in the chambers, the increment of positive units was accelerated (Figure 3d and Figure S8). 96.4% cell-containing chambers can be differentiated from negative one in 0.5 h, 1 h earlier than that of stationary cells. Similarly, significant fluctuation of the number of positive chambers appeared after that. After 3h incubation, the intensity of all the chambers containing cells was over the threshold. The fluctuation of positive counting may be a consequence of further reduction of resorufin to non-fluorescent dihydroresorufin and the inversion of this process. The similar phenomenon was also observed in 96 well plate (Figure S7d).

When we conducted the experiments with *E. coli* O114 and K-12, the similar results was obtained (Figure S9 and S10). For O114 and K-12, the first peak of positive fraction was 98.56% and 98.61% at 2 and 1.5 h, respectively. During further incubation, the positive fraction declined to 45.26 % for O114 and 38.3% for K-12. The detection of all O114 and K-12 cells were achieved till the positive fraction reached to 1 at 4.5 and 5.5 h, respectively. Since the reversible reduction of resorufin and cellular heterogeneity on metabolic activity, the first peak of positive fraction did not reach to 1 under current conditions. Despite that, more than 98% chambers with cells emitted bright fluorescence in 2 h, indicating the developed method is able to do presence/absence assay and rough quantification of cell concentrations within 2 h. Because of the ability of detecting the cells that is incapable of colony formation on agar (data shown in next part), the digital resazurin assay has equivalent performance on *E. coli* quantification in 2 h to plate counting, for which it takes over 14 h. According to the results from 96-well plate, the reduction of resorufin happened when there was over 50% LB in test solution (Figure S7d). To overcome the absorption of PDMS and supply sufficient nutrition in confined volume, we directly used the 1× LB to conduct the on-chip experiments, which may cause the fluctuation of positive counting. Therefore, it is possible to eliminate this phenomenon by sophisticated on-chip optimization of LB concentration or surface treatment to avoid excessive cellular activity.

### Quantification performance of digital resazurin assay

The quantitative performance of the developed method was evaluated using *E. coli* EK-19 in stationary phase. The samples were labelled with the diluted factors (X_dil_), for example, X_dil_ 0.05 means 20-fold dilution from stock sample. The serial solutions of EK-19 were prepared from a stock sample with 2×10^8^ CFU/mL (X_dil_1). The chamber array was filled at low λ, so there was small chance of more than one cell in a chamber. The fluorescent micrographs of the chips with different samples were captured after 3 h incubation at 42°C (Figure 4a). The number of positive chambers decreases as the dilution of *E. coli* samples. The Poisson distribution was applied to correct the cell numbers in chip with the input of the numbers of positive chambers (see Note S4 in supplemental information). The measured concentration has a good correlation with X_dil_ (R^2^ = 0.997) (Figure 4b). We also used plate counting to enumerate the *E. coli* in serial samples and observed a linear trend from the measured concentrations of cells at four diluted factors (R^2^ = 0.99, Figure S11). The results demonstrate the feasibility of digital resazurin assay for accurately quantifying the environmental *E. coli*. To further access the accuracy of this device, an EK-19 sample were simultaneously determined by the chip-based method and plate counting. As indicated in Figure 4c, the measured concentration from developed device was 1.57 times higher than that from plate counting. It was not caused by the cross-contamination of samples since the control panels had no positive signals. After careful inspection of the whole array, we found that all the positive chambers contained proliferative *E. coli* colony and they were perfectly matched in merged channel (Figure S12), showing the specificity of the reaction. In the sample panels with low *E. coli* concentrations (X_dil_ 0.05 and 0.025, λ=0.0175 and 0.0088), a few positive chambers scattered individually, demonstrating the good isolation of the chambers through the whole array. When zooming in the zone of positive chamber, merely 6 μm interval between adjacent chambers is able to offer enough bonding force to avoid the cross-talking of fluorescein and cells. Therefore, it is confirmed that the detection results of chip-based method reflect the actual cell concentrations. This phenomenon was also reported in the published paper.^43^ We speculated that the quantitative difference between our method and plate counting may result from the high sensitivity of the digital resazurin assay that the cells with low metabolic activity, which are unable to form visible colonies on agar plate, still can produce detectable fluorescent to report their presence. The smaller CV of chip-based method (4.46 %) than that of plate counting (10.51%) indicates the high reproducibility of *E. coli* quantification in 3D chamber array chip.

**Figure 4.**
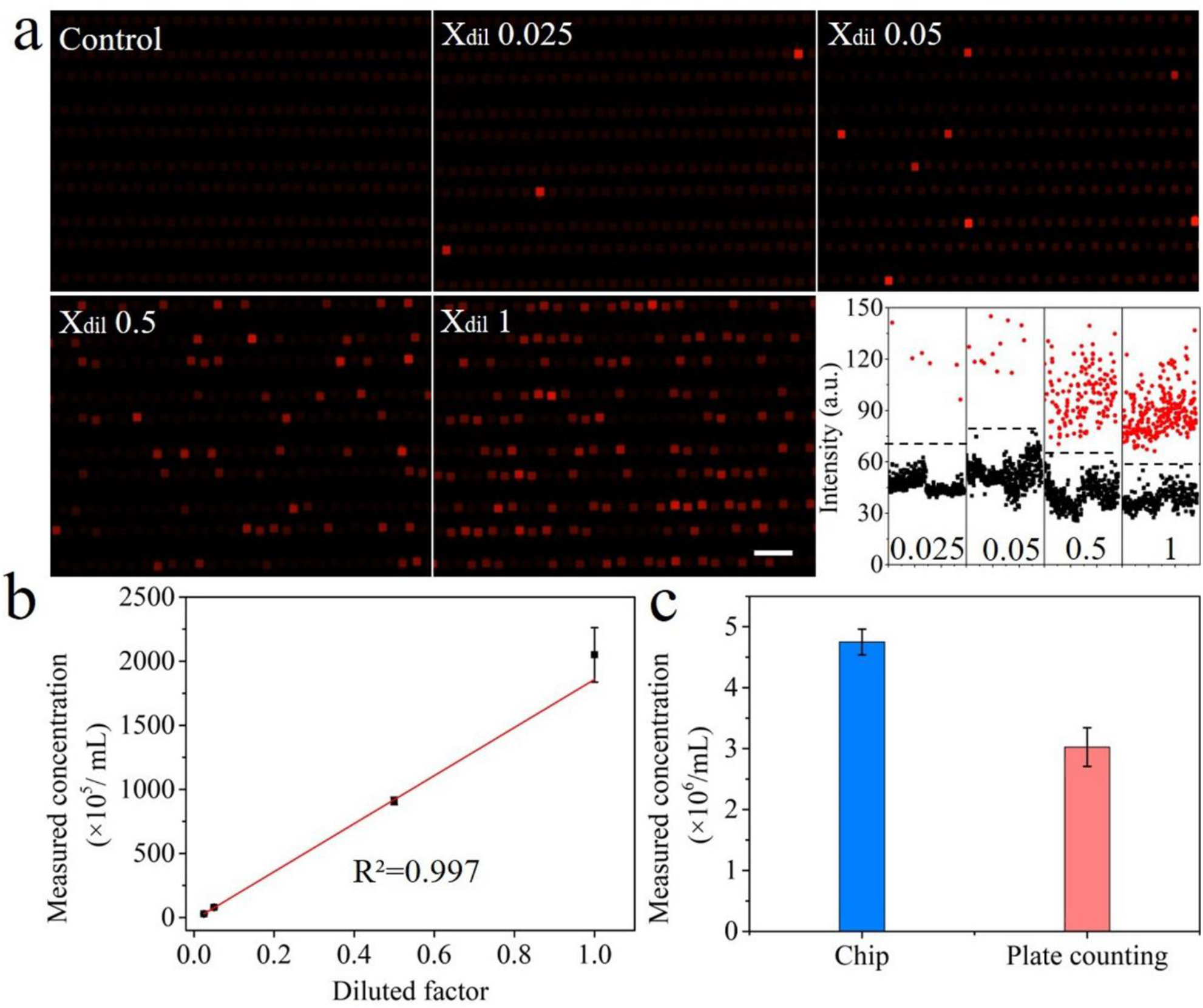
The quantification performance of digital resazurin assay in 3D chamber array. a) Fluorescent images of the 3D chamber array with various concentrations of EK-19. The threshold equals the sum of mean intensity of the negative chambers and triple standard deviation. Scale bar denotes 200 μm. b) The correlation between the measured concentrations and diluted factors. c) The quantifications of EK-19 sample using digital resazurin assay and plate counting. Error bars represent standard deviation of three replications.

### Single-cell antibiotic susceptibility testing

In consideration of the good performance of digital resazurin assay, this metabolism-dependent method was applied to conduct AST for antibiotics screening and MIC determination. Meanwhile, by capturing single *E. coli* in chamber array, the developed method allows to characterize the heterogeneity of antibiotic resistance at single-cell resolution. We firstly conducted reference testing of AST in 96-well plate (Figure S13). It was confirmed that *E. coli* O114 is susceptible to ampicillin while EK-19 and K-12 are resistant to ampicillin. The growth of these strains was inhibited in the present of polymyxin B (PMB). Subsequently, sc-AST was accomplished in developed devices by dispensing single bacteria with different concentrations of antibiotics in chambers. Enumerating the positive chambers after incubation gives the subpopulation of cells that are resistant to a certain concentration of antibiotics. The groups without antibiotics were set as controls to determinate the cell concentrations of the test samples. The cell viability is defined as the percentage of resistant cells in the populations. The AST of O114 to ampicillin was conducted in the chips with 70 μm height chambers (175 pL) and 50% LB. With the increasement of ampicillin, the cell viability of *E. coli* O114 declined gradually and all the cells were killed at 3.5 μg/mL ampicillin (Figure 5a and 5b). In contrast, the cell viability of EK-19 and K-12 was unaffected by ampicillin, which is consistent with the results in 96-well plate. Ampicillin acts as a transpeptidase inhibitor to block cell wall synthesis in binary fission, finally leading to cell lysis. The chambers with the height of 45 μm (112 pL) contain low content of ampicillin, requiring higher concentrations of ampicillin to inhibit the growth of all cells (Figure S14). Our results support that the inhibition efficiency of ampicillin is determined by the amount of antibiotic used per cell rather than the antibiotic concentration.^27^ This finding may also apply to other β-lactam antibiotics that act on cell wall synthesis. To screen effective antibiotics for the ampicillin-resistant strains, the susceptibility of EK-19 and K-12 to PMB was tested in our device. Unlike gradual decline of cell viability in the addition of ampicillin, the cell viability dropped dramatically at 2.5 μg/mL PMB for EK-19 (2.63%) and O114 (2.74%), and 1 μg/mL PMB for K-12 (7.4%). Beyond those points, the cell viability of 3 strains decreased slowly to 0 (Figure 5c). The results show the heteroresistance of *E. coli* to PMB and indicate that only a small fraction of cells (less than 8%) demonstrated an increased resistance, which may be responsible for the evolution of resistance. The PMB MICs for EK-19, O114 and K-12 were 5, 4.5 and 2 μg/mL, respectively (Figure 5d). The experiment successfully confirmed the digital resazurin assay has good functionality in antibiotics screening and the determination of the MICs for different bacterial strains, as well as quantifying the phenotype of heterogeneous resistance at single-cell level. With single-cell sensitivity and broad dynamic range, the developed platform could be a useful tool to perform comprehensive AST directly on the bacterial population in original samples for the improvement of patient outcome.

**Figure 5.**
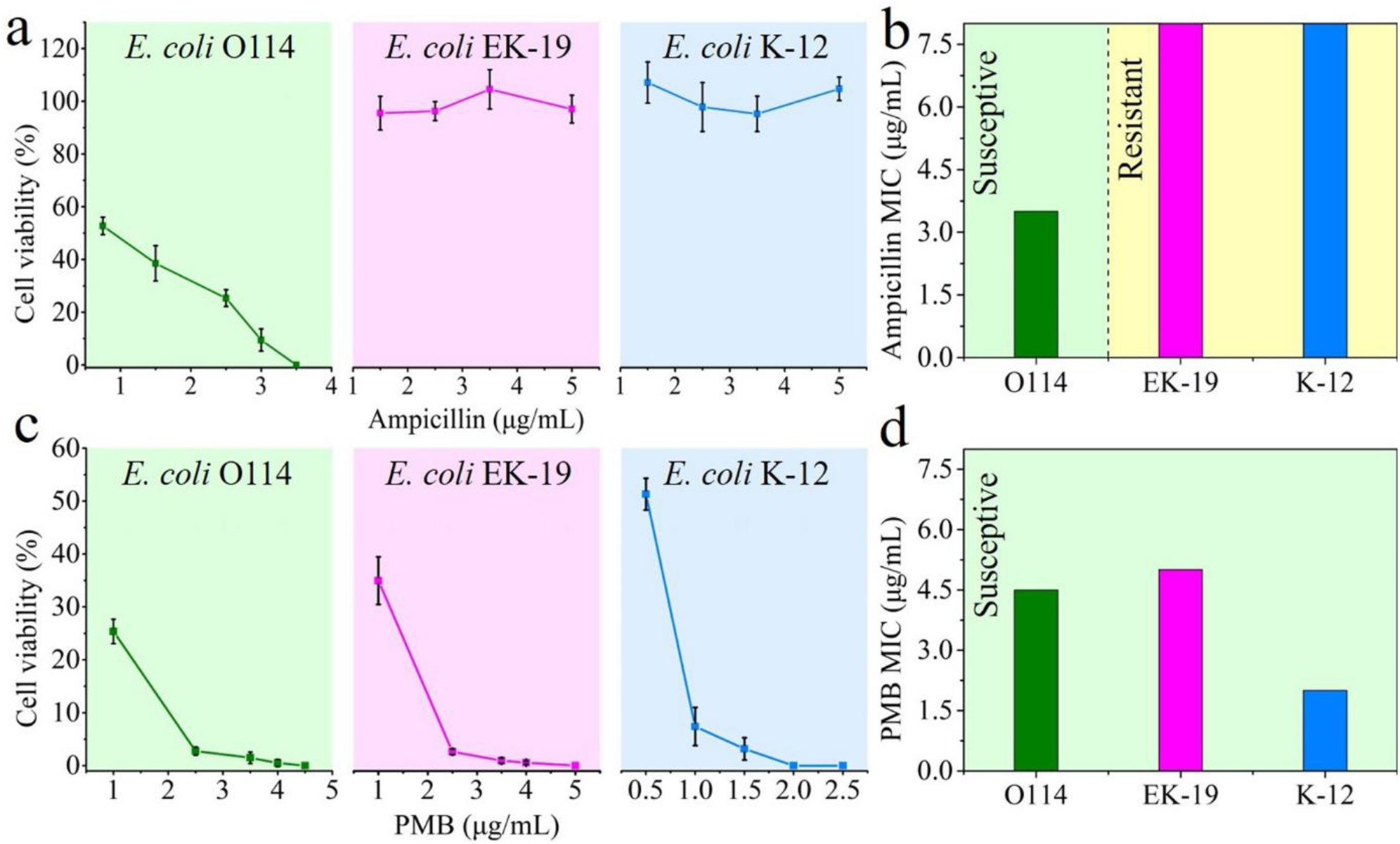
Sc-AST and MIC of *E. coli* strains treated with ampicillin and PMB. a) The Sc-AST of *E. coli* strains to ampicillin. b) MIC of *E. coli* strains exposed to ampicillin. c) The Sc-AST of *E. coli* strains to PMB. d) MIC of *E. coli* strains exposed to PMB. Error bars represent on standard deviation of three replications. The AST results of ampicillin to *E. coli* O114 were obtained by using 50% LB and the chips with 70 μm-height chambers. The others were obtained with 100% LB in the 45 μm-height chambers.

### Lyophilization and rehydration of preloaded solution

Thanks to the air permeability of PDMS, the degassed device enables automatic sample compartmentalization without the requirement of external instruments and has the possibility to perform on-site AST. In order to simplify the operation procedure, a pre-mixed solution (resazurin, medium and antibiotics) was introduced in chips and lyophilized under vacuum (Figure 6a and Video S3). With air-tight packaging, the negative pressure in chip could keep for long period and the devices with lyophilized reagents would be ready-to-use. The end users only need to load the samples into chips to reconstitute the test solution (Figure S15). After that, the filled chambers were enclosed by thermosetting oil (Figure 6a). This approach would reduce factitious error and improve the detection throughput. Due to the fast diffusion in picolitre scale, the preload reagents may leak into main channel before oil isolation, leading to the reduction of the medium, indicator, and especially for antibiotics. To overcome the leakage, we modified a 20 G needle for the introduction of sample (Figure S16). The needle with sample and oil phase was directly inserted into the inlet, which prevents the chip from losing negative pressure. The chambers can be filled by sample solution and isolated by oil sequentially in a moment (∼2 s), thus avoiding the leakage of the resolved reagents. To test the performance of needle-mediated sample introduction, a mixture containing resazurin and LB was loaded into the chamber array in advance and lyophilized. As shown in Figure 6b, the average intensity of the chambers before lyophilization was 61.89 with CV of 8.3%. The chips were rehydrated with 8 μL PBS by needle- or tip-mediated sample loading. The average intensity (69.06, CV = 10.1%) of the chambers rehydrated by using needle-mediated sample loading was slightly higher than that before lyophilization. The average intensity of the chambers rehydrated by tip-mediated sample loading reduced to 47.78 (CV = 11.05%). The lower intensity indicates the leakage of preloaded reagents and only 77.2% of them remained in the chambers. The reduction of preloaded reagents has limited effect on the detection of viable *E. coli* due to the excess of resazurin and medium (Figure 6c). In this sense, the performance of digital resazurin assay in rehydrated chip may be better than in fresh solution chip because of the higher signal/noise ratio (S/N 2.5 for the former and 1.83 for the latter). The results may differ when preloading the solutions with antibiotics for AST (Figure 6c). In the chip with fresh solution, the chambers had weak and uniform fluorescence because most of EK-19 were killed by PMB with the concentration of 2.5 μg/mL. The similar results were observed in the chips rehydrated by needle-mediated sample loading, indicating the reconstituted PMB in chambers was able to inhibit the growth of EK-19 cells. But there were some chambers with increased intensity in the chips rehydrated by needle-mediated sample loading. This phenomenon results from the decreased concentration of PMB during tip-mediated rehydration. The experiments indicate that the modified needle enable to avoid the leakage of preloaded reagents during rehydration. In combination of lyophilization, the self-powered chip has potential for fast and on-site sc-AST for drug selection and MIC determination. The required instruments are a small incubator for resazurin reaction and a common fluorescence microscope for chip inspection, which are essential facilities in most clinics (Figure S17). As reported in published papers,^44, 45^ several miniaturized apparatuses with the integration of incubator and fluorescence detector were developed for nucleic acid detection. They would further improve the portability of the sc-AST in developed chip.

**Figure 6.**
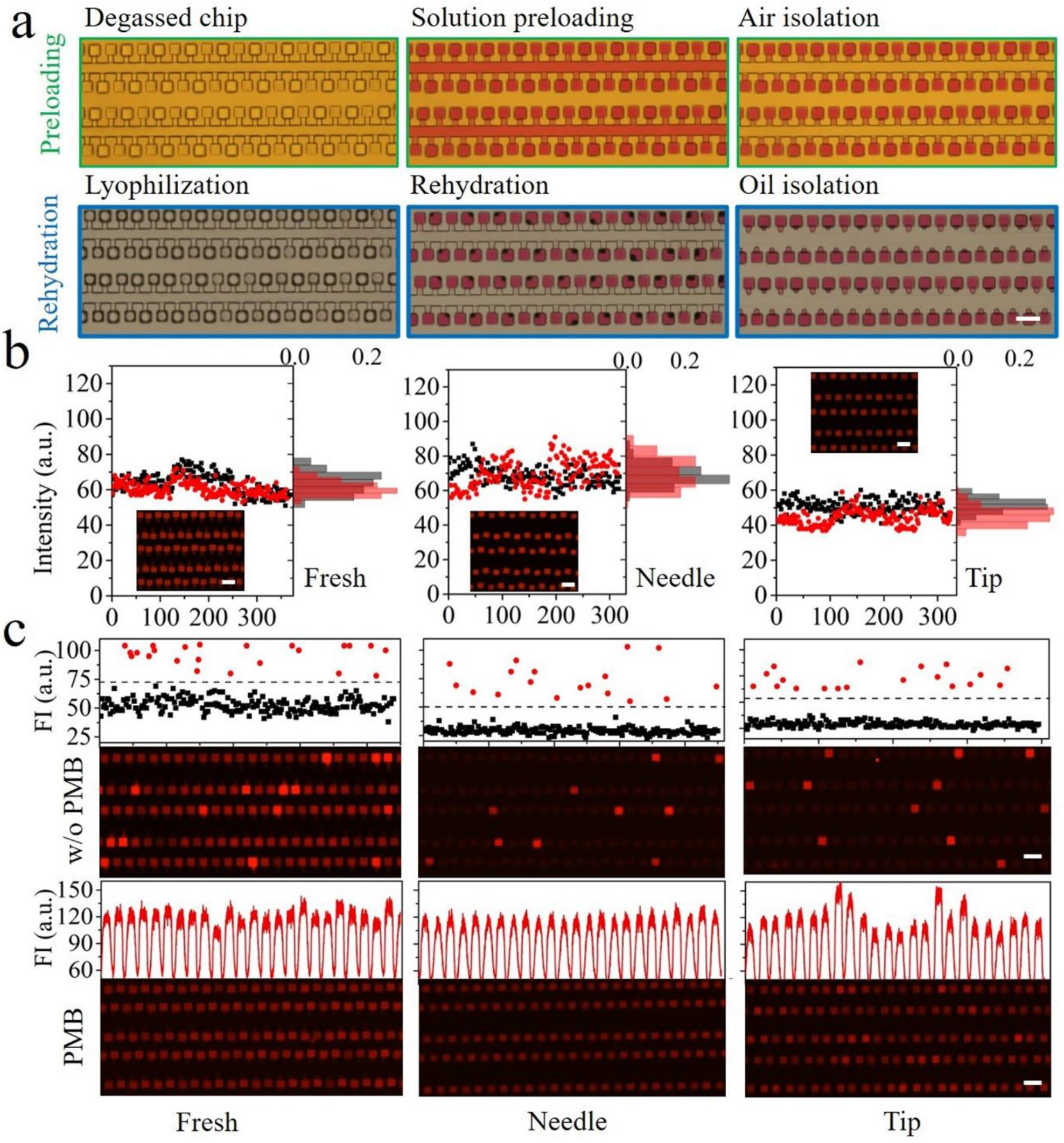
Needle-mediated rehydration of preloaded reagents. a) Lyophilization and rehydration of preloaded solution for on-site AST. b) The rehydration of chips by needle- or tip-mediated sample loading. The black dots represent the fluorescence intensity of the chambers next to the inlet. The red dots represent the fluorescence intensity of the chambers next to the outlets. c) The performance of needle-mediated rehydration in *E. coli* detection and AST. The chips with lyophilized reagents were used after degassing without airtight packing. In the experiments, *E. coli* EK-19 at 2.4 × 10^7^ CFU/mL and 2.5 mg/mL PMB were used. Scale bars denote 100 μm. FI: Fluorescence intensity.

## CONCLUSIONS

In summary, we provided a portable and user-friendly PDMS chip to realize rapid and digital quantification of *E. coli* and demonstrated its application in sc-AST assay. The developed device has two distinctive characters: 1) capillary valve-based flow distributor, 2) high-density 3D chamber array. The capillary valve-based flow distributer realized flow equidstribution for scale-up of 64 parallel channels. The high density of reaction units and wide dynamic range were achieved by the 3D arrangement of chamber array in developed chip. Driven by the negative pressure in degassed PDMS, the sample solution can be automatically distributed to the chambers through the bidirectional network in 2 s. We developed a digital resazurin assay in 3D chamber array to realize single *E. coli* quantification in 2 h. The digital resazurin assay are sensitive enough to detect the low active *E. coli* and has favorable quantification performance in compared to plate counting. The sc-AST for antibiotics screening and determination of MIC was achieved in the developed chip. The experiments also evaluated the heterogenetic resistance to antibiotics at single-cell resolution. Moreover, we investigated the preload of test reagents to simply the handling process at user side and realized the leakage-free rehydration of dried reagents by using the needle-mediated sample loading. The integrated technologies of self-powered microfluidic chip, digital resazurin assay and lyophilization have potential to conduct on-site AST assay to facilitate the treatment of bacterial infections and could benefit understanding the mechanisms and evolution of antibiotic resistance.

## METHODS

### Chip fabrication

The chips used in this work were prepared by soft-lithography and PDMS molding (Figure S18, See supporting information).

### Bacteria culture and sample preparation

*Escherichia coli* O114 (ATCC 25922), *Escherichia coli* K-12 (ATCC 19020) and *Escherichia coli* EK-19 were inoculated in sterile Luria Broth (LB, L3522, Sigma) and incubated at 37 °C overnight with a shaking speed of 200 rpm. *E. coli* EK-19, an environmental strain, was isolated from soil in Malaysia and kindly provided by Prof. Eric P. H. Yap (Lee Kong Chian School of Medicine, Nanyang Technological University). If not mentioned, all the bacterial samples used in this work were harvested after overnight cultivation (stationary phase). The logarithmic phase *E. coli* EK-19 was obtained at the OD value of 0.15. To prepare the test solution, 1 mL *E. coli* suspension was transferred into 1.5 mL tube and centrifuged at 1540 RCF for 15 min. The supernatant was discarded and same amount of PBS (PB0344, Vivantis) was pipetted into tube to resuspend the pellet. After diluted to desired concentration with fresh PBS, the *E. coli* samples were ready for usage.

### Test solutions and optimizing test conditions in 96-well plate

The concentration of mediums, LB, Tryptic Soy Broth (T8907, Sigma), Tryptic Soy Broth without dextrose (T3938, Sigma) and Hi-Def Azure Media with 1% Glucose (3H5000, Teknova) were optimized. The optimization of resazurin (0.01-0.2 mM) in test solution and the incubation temperature (35-42°C) were also conducted. For the optimization in 96-well plate, each 320 μL test solutions consisted of 6.4 μL *E. coli* sample (∼8×10^5^ CFU/mL), 11.2 μL resazurin and 302.4 μL medium and were aliquoted into 3 parallel wells with 100 μL per well. To confirm the AST, 32 μL ampicillin (Roche) or PMB (P1004, Sigma) with different concentrations were added in above solutions. For *E. coli* detection in chip, the test solutions were composed of 6.4 μL *E. coli* sample, 11.2 μL resazurin (2 mM) and 302.4 μL LB Broth (1×) or 299.2 μL LB Broth (1×) and 3.2 μL antibiotic solutions.

### Lyophilization and rehydration of pre-mixed solution

The pre-mixed solution consisted of 6.4 μL DI water, 11.2 μL resazurin (2mM) and 302.4 μL LB Broth (1×, without NaCl) or 299.2 μL LB Broth (1×, without NaCl) and 3.2 μL antibiotic solutions. The procedures of lyophilization were similar to test sample loading (Figure S17). To perform AST assay, a modified needle (20G) with 8 μL sample solution and 20 μL oil was used to puncture the tape and inserted into inlet to rehydrate the preloaded reagents. One important note in rehydration is keeping osmotic pressure of rehydrated solution fit with that of *E. coli*. If standard LB existed in pre-mixed solution, the *E. coli* samples in DI water should be used for rehydration.

### Data Acquisition and analysis

All the fluorescent micrographs were acquired by Zeiss microscope (Axio Observer 7, Germany). After incubation, the chips were observed by using 10× lens (0.3 N.A.) and captured by monochrome camera (Axiocam 705, Zeiss) under PHY channel (546/10 nm; 585/40 nm). The EGFP channel (480nm/30 nm; 535/40 nm) was applied to visualize the fluorescence beads (PS, 2 μm). The time-lapse imaging of *E. coli* detection course was realized with the help of Time-lapse module of Zen Pro (Version 3.3). The common processing of micrographs including addition of pseudo color, adjustment of brightness, subtraction of background, stitching the images of multiple views, and merging channels was also done in Zen Pro. The bright-field videos showing hydrodynamic behaviors were obtained by OLYMPUS microscope (IX81, Japan) equipped with a colorful camera (NIKON). The correlation analysis and T-test were conducted by using Fitting and two samples t-test functions of OriginPro 9.0. The reduction reactions of resazurin in 96-well plate were monitored by plate reader (SPARK 10M, TECAN, Switzerland) under 560/590 nm. The optical density of test solution without resazurin at 600 nm was recorded to indicate the cell growth during incubation.

## Supporting information

Supplemental methods and results

## ASSOCIATED CONTENT

### Supporting Information

The following files are available free of charge.

Additional details regarding the limitation of 2D chamber array (Figure S1), chip fabrication and characterization (Note S2, S3, S4, S6 and Figure S2, S4, S5, S6, S18), the fluorescence images of *E. coli* detection (Figure S8, S9, S10, S12), the verification of the results by conventional method (Figure S7, S11, S13), the relationship between the inhibition efficiency and the amount of antibiotics (Figure S14) and the workflow of chip operation and on-site AST (Note S1 and Figure S3, S15, S16, S17) (PDF)

Supporting videos showing the flow distributor, automatic sample loading and rehydration (Video S1, S2, S3) (MP4).

## AUTHOR INFORMATION

### Corresponding Author

*.

### Author Contributions

The manuscript was written through contributions of all authors. All authors have given approval to the final version of the manuscript. §These authors contributed equally.

### Notes

The authors declare no competing financial interest.

## ACKNOWLEDGMENT

The authors thank Prof. Eric P. H. Yap (Lee Kong Chian School of Medicine, Nanyang Technological University) for kindly providing an environmental *E. coli* strain (EK-19).

## Table of Content

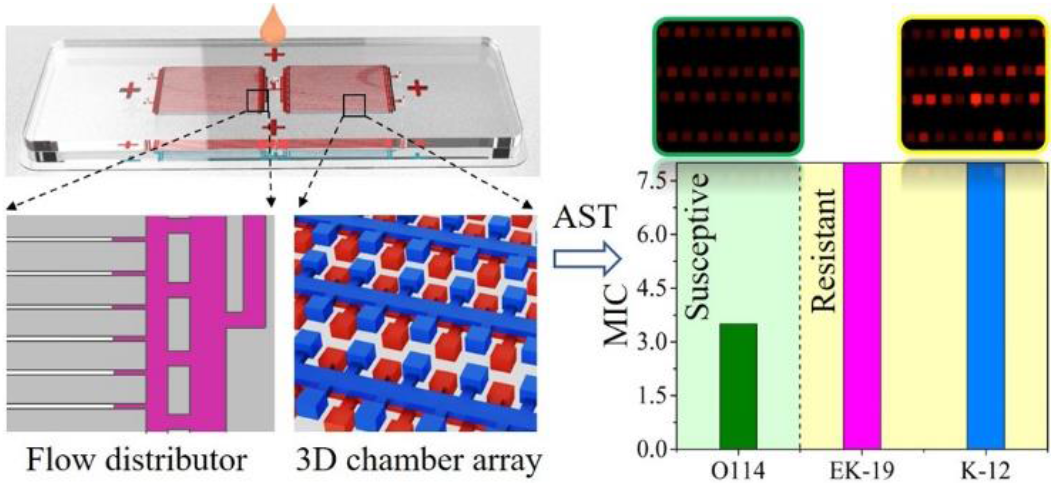

