## Supplemental methods and results for "A portable and high-integrated 3D microfluidic chip for bacterial quantification and antibiotic susceptibility testing"

**Content**

**Figure S1**. The limitation on chamber integration in 2D chamber array.
**Figure S2**. The anti-evaporation performance of ultra-thin chip.
**Figure S3**. Flow chart showing chip operation and self-driven sample loading.
**Figure S4**. The size of chambers.
**Figure S5**. The comparison between the intensity of chambers in upper and lower PDMS.
**Figure S6**. The simulation on the bead distribution in 3D chamber array.
**Figure S7.** The optimization of resazurin, medium and reaction temperature in 96 well plate.
**Figure S8.** Time-lapse imaging of log phase *E. coli* EK-19 detection.
**Figure S9.** Time-lapse imaging of *E. coli* K-12 detection.
**Figure S10**. Time-lapse imaging of *E. coli* O114 detection.
**Figure S11**. Enumerating the *E. coli* on agar plate.
**Figure S12**. The confirmation of the positive chambers.
**Figure S13**. AST in 96 well plate.
**Figure S14**. The inhibition efficiency of ampicillin to *E. coli* O114 in 45 and 70 µm chambers.
**Figure S15**. Workflow of reagent preloading and on-site AST.
**Figure S16**. Liquid transferring and needle-mediated sample loading.
**Figure S17**. The instruments required in on-site AST.
**Figure S18**. Flow chart of chip fabrication.
**Video S1.** Flow distributor.
**Video S2.** Self-driven sample loading and oil isolation.
**Video S3.** Sample preloading and rehydration

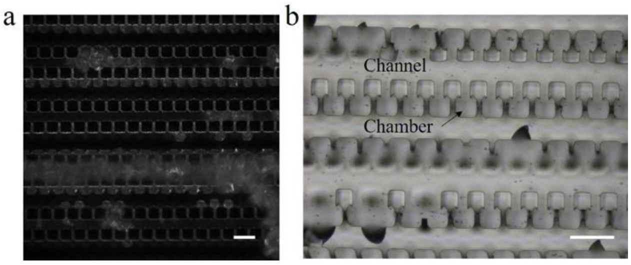


**Figure S1**. The limitation on chamber integration in 2D chamber array. a) The mold with 20 μm interval after PDMS releasing. B) The broken chambers in PDMS chip. When the interval between chambers decreases to 20 μm, it is a challenge to total remove the unsolidified photoresist. And the mechanical strength of 20 μm PDMS interval cannot resist the frictional drag during peeling off the PDMS elastomer and it tends to fracture, causing chamber connection. Scale bar denotes 100 μm.

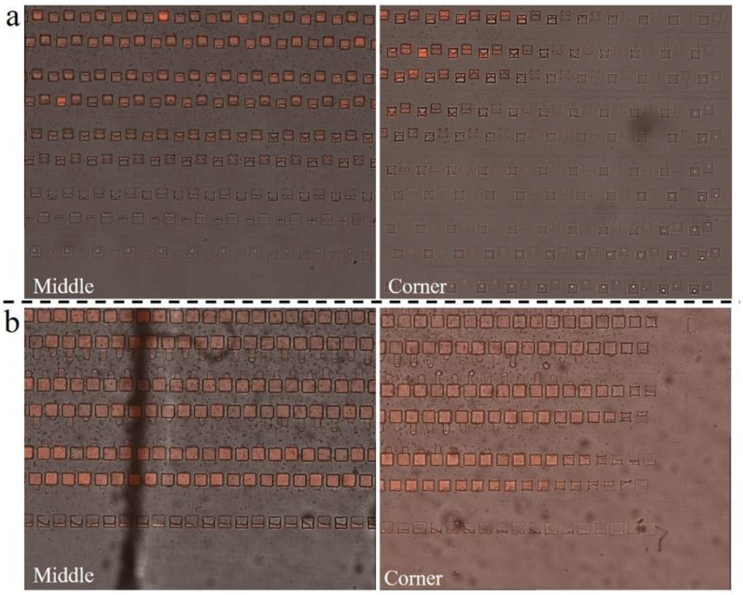


**Figure S2**. The anti-evaporation performance of ultra-thin chip. a) the water loss of peripheric chambers in normal chip after 42 ℃ for 3h. Left: the middle of chamber array; Right: the corner of chamber array. b) the water loss of peripheric chambers in ultra-thin chip after 42 ℃ for 3h. Left: the middle of chamber array; Right: the corner of chamber array.

**Note S1: Chip operation**

Before sample loading, the microfluidic chip was firstly degassed in a vacuum chamber for 50 min to produce negative pressure in chip (Figure S3). After that, the cover tape on inlet was sticked by a needle and the tip with 10 μL test solution and oil phase was inserted in the inlet. The oil phase containing 0.65g 5 cSt silicone oil (317667, Sigma) with 0.12 mM platinum catalyst (479527, Sigma), 0.25g 50 cSt silicone oil (378356, Sigma), 0.15g PDMS monomers (RTV-615 A) and 0.05g PDMS cross-linker (RTV-615 B) was on the top of test solution to isolate the chambers. When the oil phase entered into the outlets, the PDMS batteries and cover tape were removed followed by sealing the inlet and outlets with 10:1 PDMS prepolymer. An optional procedure, keeping the chip in air and normal temperature for ~15 min, helps to release the negative pressure in PDMS and reduce the water evaporation of peripheral chambers. Finally, a coverslip was put on the surface of PDMS to be a vapor barrier and the enclosed chip was incubated at 42 °C in a miniature incubator (Okolab, Italy).

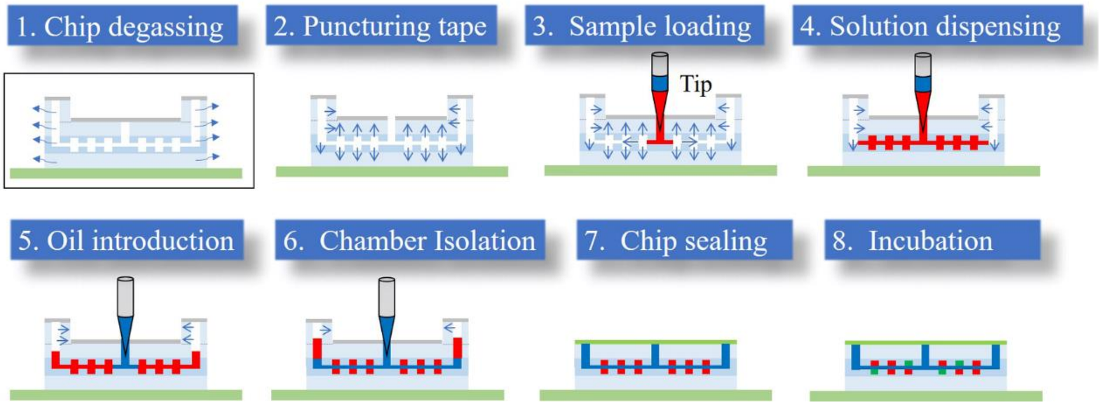


**Figure S3**. Flow chart showing chip operation and self-driven sample loading. The air permeability of PDMS contributes to the self-driven liquid distribution. After the air in PDMS was evacuated by vacuum pump, the negative pressure was produced and stored in PDMS bulk, providing the power for liquid flowing along the channel network to the whole chamber array.

**Note S2: The measurement of chamber size**

The height of chambers and channels was determined by a step profiler using SU-8 mold. The rising height of pin from wafer to the top surface of photoresist micropillars denoted the height of microwell. To measure the chamber area, the chip was filled with the liquid of red colouring and then captured by camera under inverted microscope (Olympus, Japan). Image J was used for analyzing and extracting the size information of red spots. The height of chamber multiplied by the average area of chambers equals the volume of microwell. The height profile of chamber array was plotted in Figure S4a. Each peak represents one chamber and the peak value was extracted as the height of it. The average height of chamber was 45.2 μm with variable coefficient (CV) of 1.76%. The average area of chambers was acquired via analyzing the micrograph of chambers filled by red colouring in Image J. The distribution of chamber area was narrow with a CV value of 3.72%, as shown in Figure, and the mean value was 2469 $\mu m^2$.

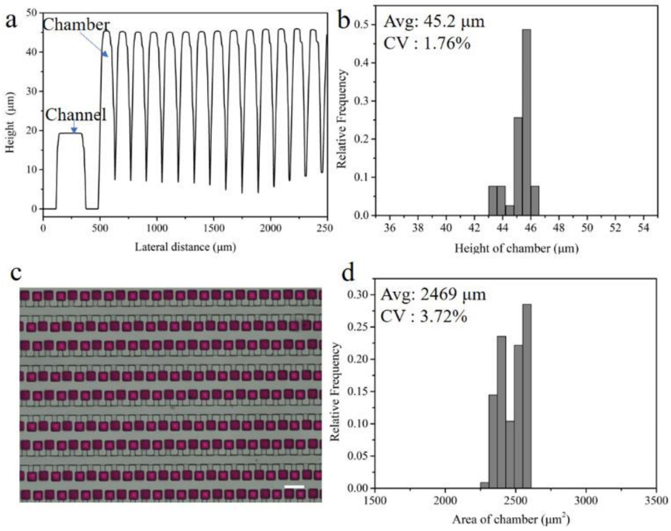


**Figure S4**. The size of chambers. a) The profile of the photoresist structures on wafer. b) The distribution of chamber height. c) A picture of 3D chamber array for chamber area measurement. Scale bar denotes 100 μm. d) The distribution of chamber area.

**Note S3: The lens for uniform fluorescence detection of 3D chamber array**

The 3D chamber array of the developed device consists of two 2D arrays that are in two planes. Taking the bonding surface of the two PDMS elastomers as a reference, the height range of the 3D chamber array is from -45.2 to +45.2 μm with a span of 90.4 μm. To get a clear and homogeneous fluorescence imaging of the 3D chamber array at one snap, the depth of light collecting of the lens is a key parameter, which relates to the depth of field (DOF) and numerical aperture (N.A.). A 5× lens (N.A. 0.16, DOF 11.4 μm) was applied to acquire the fluorescence pictures of the 3D array filled with resorufin solution. Although it has ability of visualizing all chambers with highly uniform intensity (data not shown), the lens collects the light coming at a small angle (18.4°), leading to low sensitivity of fluorescence detection. A lens with higher N.A. can improve the sensitivity of fluorescence detection through collecting more light coming at a bigger angle but has decreased DOF and the depth of light collecting, which may cause the uneven fluorescence imaging. Therefore, a 10× magnification lens (0.3 N.A. DOF 3.1 μm) was applied and can collecting light coming at an angle of 34.9°.

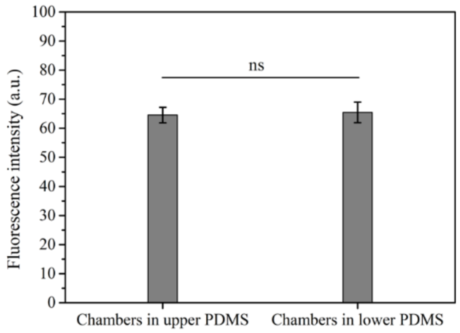


**Figure S5**. The comparison between the intensity of chambers in upper and lower PDMS (*P>0.05*).

**Note S4: Simulation on the bead distribution in 3D chamber array**

As indicated in Figure 2i, the number of beads in upper chambers was slightly bigger than that in lower. In order to figure out the reasons, we used the COMSOL to illustrate the bead encapsulation in the 3D chamber array. The model of chamber array and channel in COMSOL was built as Figure S6a, which had the same size and chamber arrangement as that in chip. Because of the porosity of PDMS, negative pressure in PDMS is generated after degassing and serves as the power source driving the liquid and cells into chambers and keeping them in chambers. It is a big challenge to simulate the liquid flow and *E. coli* movement driven by negative pressure in porous PDMS using COMSOL. Therefore, the branch channel in the model was removed to facilitate beads capture and the beads were released at the speed of 5 mm/s from inlet. Firstly, the velocity of liquid at the entry of chambers was investigated. There was no significant difference between the liquid velocity at the entry of upper and lower chambers (Figure S6b). The result indicates that the beads moving in main channel have same chance to be captured by upper or lower chambers. We recorded the number of beads entering into upper and lower chambers at a second as a surrogate of counting the beads captured in upper and lower chambers of degassed chip to investigate the probability of beads distribution in two layers, which was indicated by the ratio between the number of beads in upper and lower chambers. When the ratio is close to 1, it is demonstrated that the distribution of beads in the upper and lower chambers is random. As shown in Figure S6c, the ratio was getting close to 1 with the increasement of the release frequency of beads and the standard deviation gradually decreased. These results confirmed that there is no obvious difference on bead distribution between upper and lower chamber array and the configuration of 3D chamber array ensures the random cell dispensing in principle.

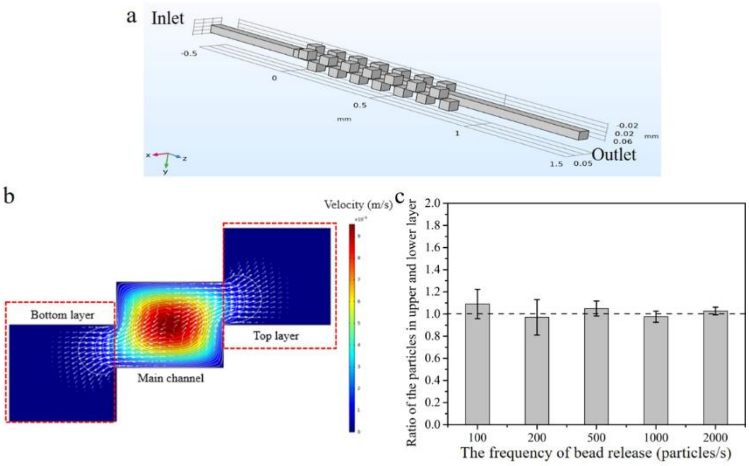


**Figure S6**. The simulation on the bead distribution in 3D chamber array. a) The simplified model of 3D chamber array in COMSOL. b) The velocity profile. c) The ratio between the number of bead in upper and lower chambers.

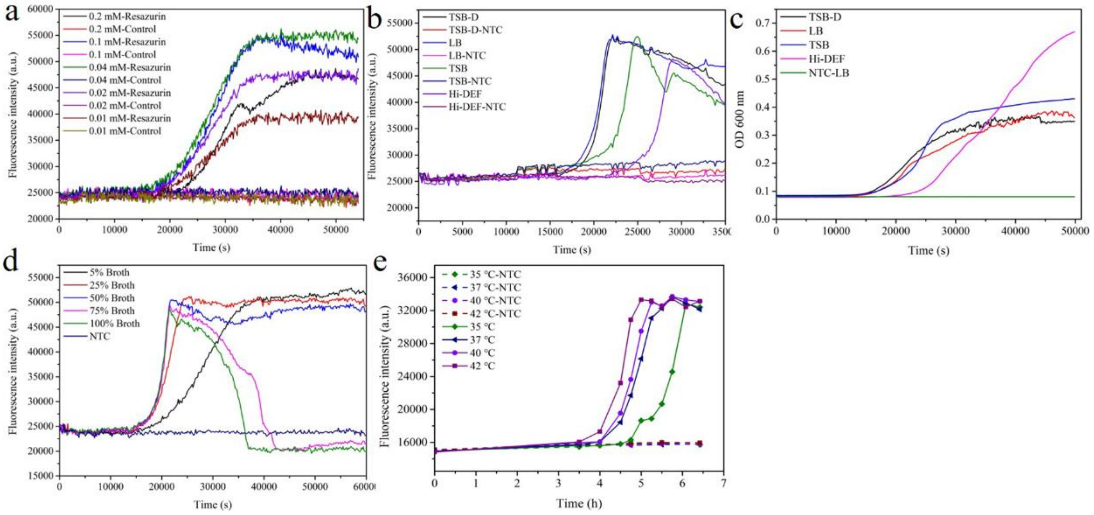


**Figure S7.** The optimization of resazurin (a), medium (b, c and d) and reaction temperature (e) for fast resazurin reaction in 96 well plate. The curves indicate the average of triplicate wells.

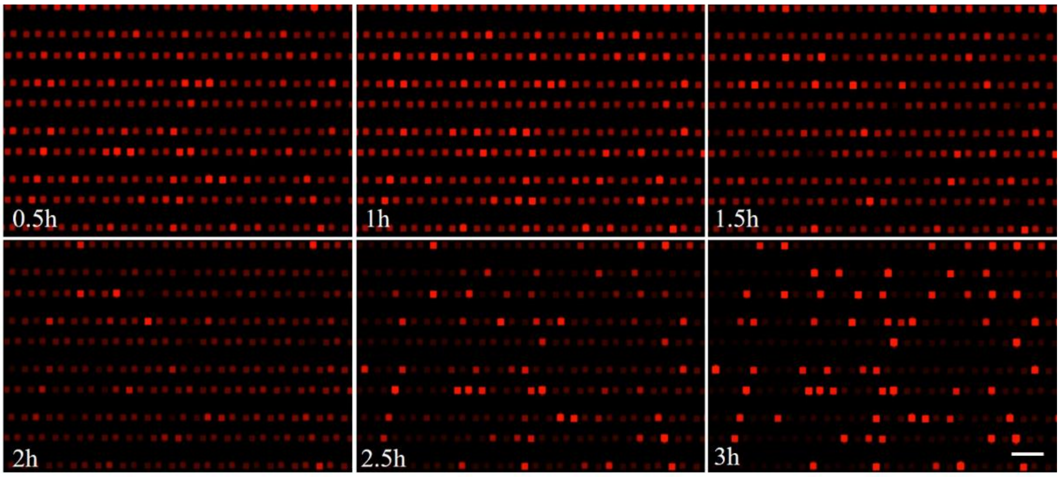


**Figure S8.** Time-lapse imaging of log phase *E. coli* EK-19 detection. The EK-19 was harvested at the OD of 0.15. Scale bar denotes 200 μm.

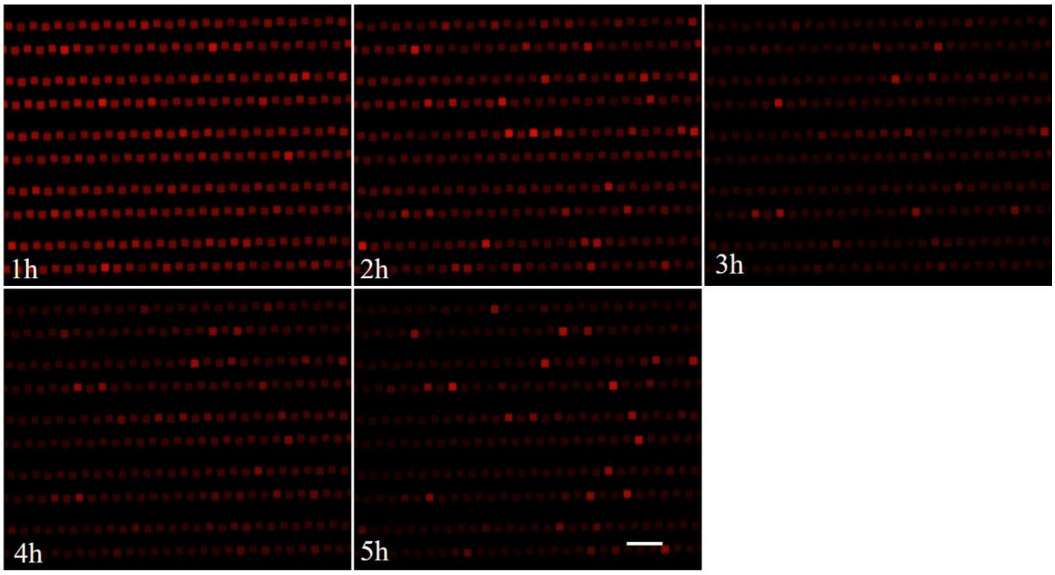


**Figure S9.** Time-lapse imaging of *E. coli* K-12 detection. Scale bar denotes 200 μm.

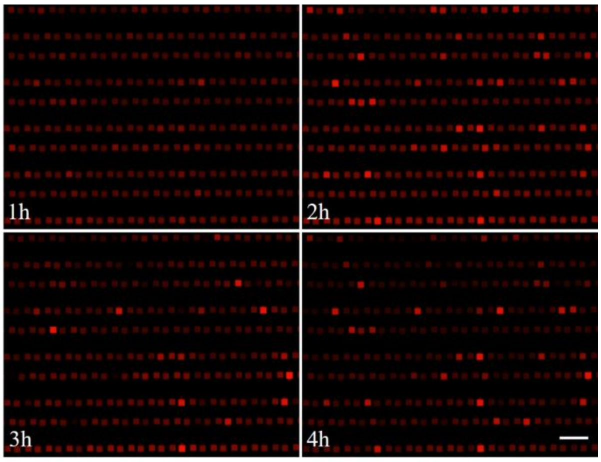

**Figure S10**. Time-lapse imaging of *E. coli* O114 detection. Scale bar denotes 200 μm.

**Note S5: The determination of *E. coli* concentration according to Poisson distribution**

Similar to digital PCR, the quantification of *E. coli* in samples depends on the random distribution of single cell in chamber array. It has been demonstrated that the cells were randomly encapsuled in chambers and the number of cells in each chambers fitted Poisson distribution. Due to the high chance for multiple cells in one chamber at a high initial concentration, the correction of targets number from the number of positive chambers in chip was necessary by using Poisson distribution formula:

$$P(n, \lambda) = (\lambda^n \cdot e^{-\lambda})/n! \quad (1)$$

n denotes the cell number in a chamber (0, 1, 2, 3…) and $\lambda$ is the average number of cells per chamber (total number of *E. coli* in chip / the number of chambers). P is the probability that the chamber contains n cells. The stochastic distribution of targets gives the chamber two conditions, with cells and without cells. When a chamber shows bright fluorescence after incubation, there is at least 1 cell in it and the probability of that is

$$P(n > 0) = 1 - P(n = 0) = 1 - e^{-\lambda} \quad (2)$$

And, it also equals the proportion of positive chambers in the chip, then

$$P(n > 0) = 1 - e^{-\lambda} = a/b \quad (3)$$

In which *a* and *b* are the number of positive chambers and the total amount of chambers per chip, respectively. After incubation, the counting of positive is a constant. Therefore, the $\lambda$ can be calculated as follow:

$$\lambda = -ln(1 - a/b) \quad (4)$$

As indicated above, $\lambda$ equals the number of *E. coli* in chip divided by the number of chambers. The cell number in chip is unknown and defined as *X*. The formula 4 can be rewritten as

$$X/b = -ln(1 - a/b) \quad (5)$$

Or

$$X = ln(1 - a/b) \cdot (-b) \quad (6)$$

In this work, the total number of chambers per chip is 22660.

So, the relations between cell number (X) and the number of positive chamber (a) are as followed:

$$X = ln(1 - a/22660) \cdot (-22660) \quad (7)$$

The volume of the chamber array is 2.55 μL. Eventually, the concentration of the *E. coli* in test solution can be calculated.

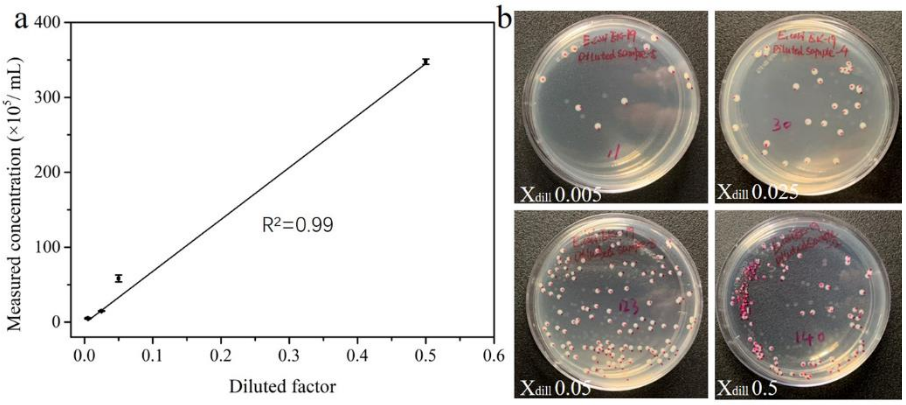


**Figure S11**. Enumerating the *E. coli* on agar plate. a) The correlation between the measured concentrations and diluted factors. b) The pictures of the agar plates after overnight incubation. Before spreading, the samples with diluted factors of 0.05, 0.025 and 0.005 were further diluted with 1000 times volume of PBS and the sample with diluted factors of 0.5 was diluted with 5000 times volume of PBS. Then, 20 μL diluted solutions was pipetted on agar plate for spreading. Error bars represent on standard deviation of three replications.

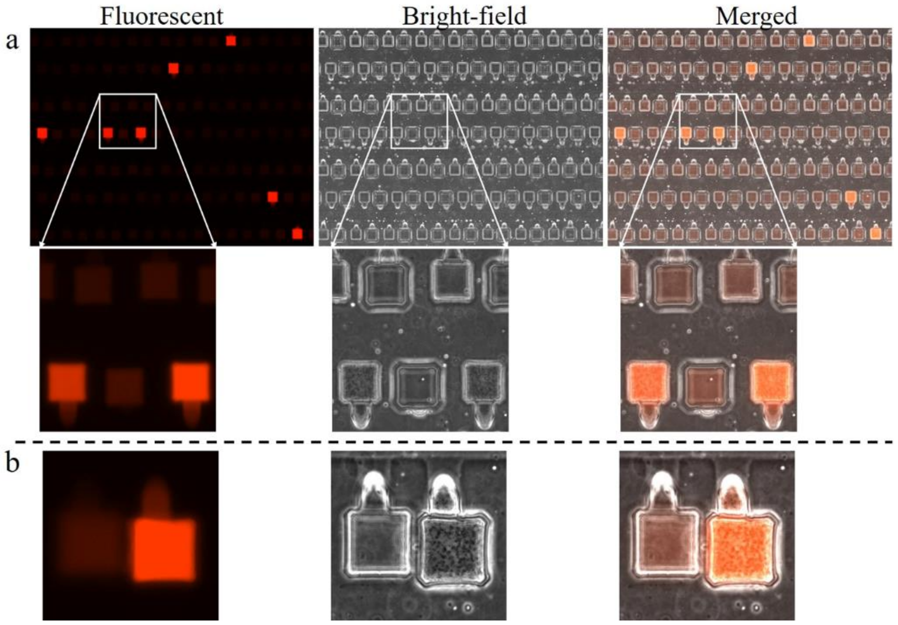


**Figure S12**. The confirmation of the positive chambers. a) The correlation between bright fluorescence and the presence of *E. coli* colonies proliferated from single cells in chambers. b) The reliable chip bonding and oil isolation.

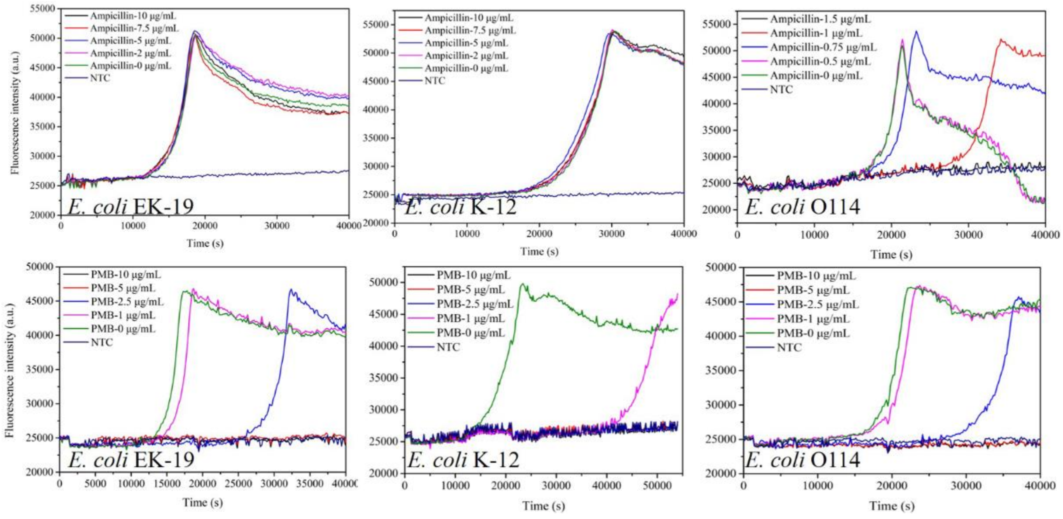


**Figure S13**. AST in 96 well plate. The curves indicate the average of triplicate wells.

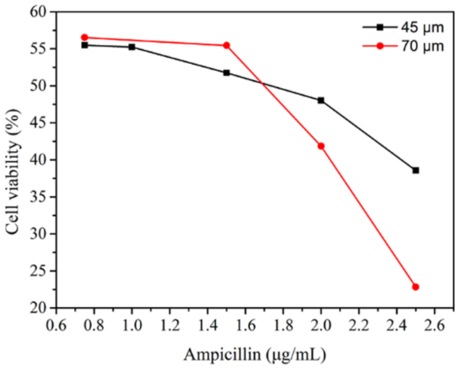


**Figure S14**. The inhibition efficiency of ampicillin to *E. coli* O114 in 45 and 70 μm chambers. 100% LB was used in this experiment.

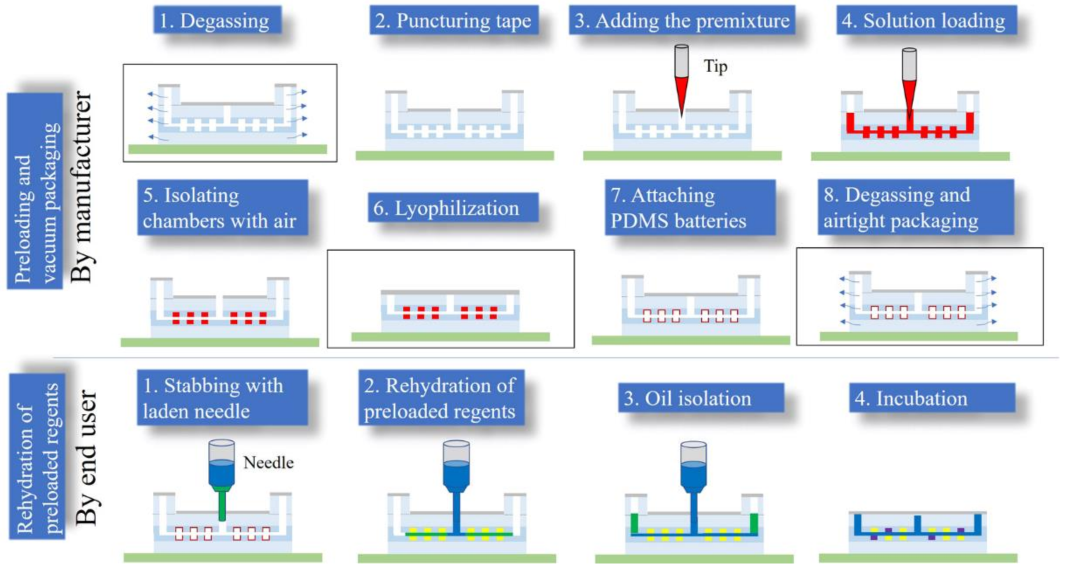


**Figure S15**. Workflow of reagent preloading and on-site AST. Reagent preloading: 1. Degassing the chip in vacuum chamber for 30 min; 2. Punching the seal tape on the inlet; 3. Adding 20 μL pre-mixed solution by using tips; 4. Pumping the pre-mixed solution into chip by negative pressure; 5. Extracting the liquid in channel network by syringe; 6. Lyophilizing the solution in a vacuum chamber; 7. Attaching PDMS batteries on the outlets of dried chip and covering the surface using seal tape; 8. Degassing the chip and vacuum packaging. Rehydration and incubation: 1. Stabbing the seal tap with laden needle; 2. Rehydrating the dried reagents in chip; 3. Isolating the chambers by oil; 4. Incubating the chips at 42 °C.

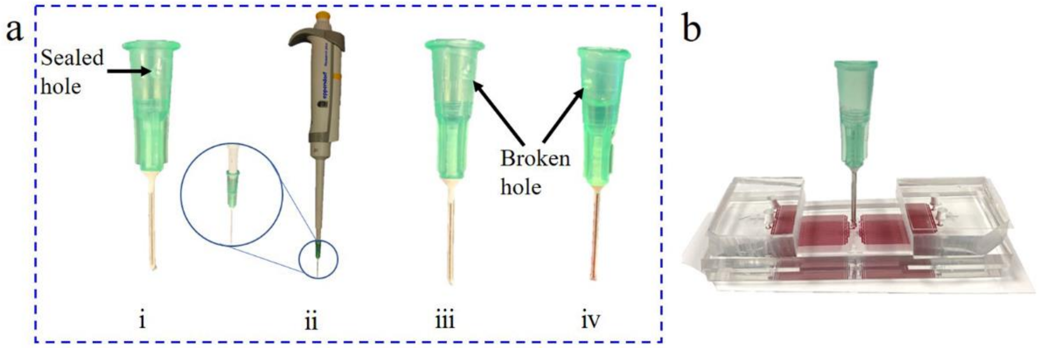


**Figure S16**. Liquid transferring (a) and needle-mediated sample loading (b). A 20 G needle (OD 0.8mm, ID 0.3mm) was cut and drilled to make a hole on its adaptor (i). The hole was used for balancing the pressure when taking off the needle from pipette. Before liquid transferring, the hole was sealed by tape. 20 μL pipette was used for transferring 8 μL sample solution into the needle, which was stored in the metal capillary (ii). Then, the tape was sticked and the needle was detached from pipette (iii). Sequentially, oil phase was added in the needle adaptor (iv). With the sharp tip, the needle can puncture the seal tape on the inlet and insert into the inlet directly.

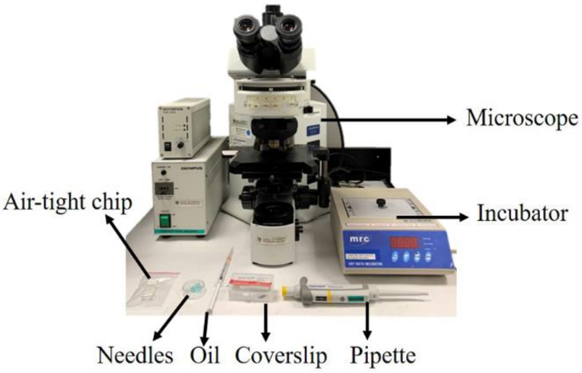


**Figure S17**. The instruments required in on-site AST.

**Note S6: Chip fabrication**

In brief, the patterns containing channel network and chamber array were designed in L-Edit v15.0 (Tanner EDA, Monrovia, CA) and transferred onto two 7-inch chrome-masks by laser writer (Nanoinprint, Singapore). For mold fabrication, the SU-8 3010 (KAYAKU, Westborough, USA) was firstly coated onto a clean silicon wafer (6 inch) at 1000 rpm for 30s to make channel layer (~20 μm). After 10 min prebaking, the wafer was patterned by using channel mask under 365 nm UV light (26.5 mW/cm2, 10s) and baked at 95℃ for 5 min to make photoresist solidify. Next, the second layer of SU-8 3010 was spun onto channel layer at the same condition. The mask of chamber array was aligned with the channel structures by a mask aligner (Karl Suss MA-6, Süss Microtec Se, Germany) and the permanent structures was formed after UV exposure (15s) and post-baking (95℃,8 min). After development and hard baking, the SU-8 structure was covered by trimethylchlorosilane solution (Sigma-Aldrich, Germany) for 10 min to facilitate PDMS release.

For thick PDMS chip fabrication (5.5mm), 5g PDMS precursor with 1g cross-linker (RTV-615, Momentive) was fully mixed, poured onto the mold and vacuumed for 15 min to remove air bubbles. The wafer was spun at 1500 rpm for 20 s and put in 80 °C oven for 3 min to make PDMS harden. Then, 44g degassed 10:1 PDMS was cast on the first PDMS layer and polymerized at 80 °C for 30 min. The PDMS replicas were peeled off and punched by 1mm puncher (WPI, USA) for the inlets and outlets. The upper and lower layers were attached on glass slides with the patterns face up. Subsequently, these two layers were put face to face and aligned manually under a stereomicroscope (Olympus,4 × lens, Japan), during which DI water could be optionally sprayed on the patterns as lubricant to facilitate the alignment and allow to repeat the alignment. The combinations were dried at 55°C overnight and exposed to UV for 2h after the removal of top glass. Afterward, the assembled devices were baked at 80 °C for over 10 h to enhance the bonding strength. For thin PDMS chip fabrication, a modified protocol was developed, which is similar to that of the thick chip fabrication. Following spin-coating of 5:1 and 10:1 PDMS sheet on the molds at 1500 rpm and 300 rpm, respectively, oxygen plasma (PDC-002, Harrick Plasma, USA) was used for bonding it with the glass slides (~500 µm) with/without holes (1.0 mm in diameter). To facilitate the releasing of glass bonded PDMS sheets, the PDMS film was firstly cut into pieces along the glass edge and then covered by ethanol for 30 min. PDMS film was punched for inlet and outlets after it was detached from wafer. The following procedures were same as the above. In the both cases of thin and thick PDMS chip, two 4 mm-thick PDMS elastomers with holes (1.5 mm in diameter) were mounted at the outlets to speed up the chamber isolation. The top surface of the assembled device was sealed with 96-well plate seal tape (Sigma-Aldrich, Germany) before degassing.

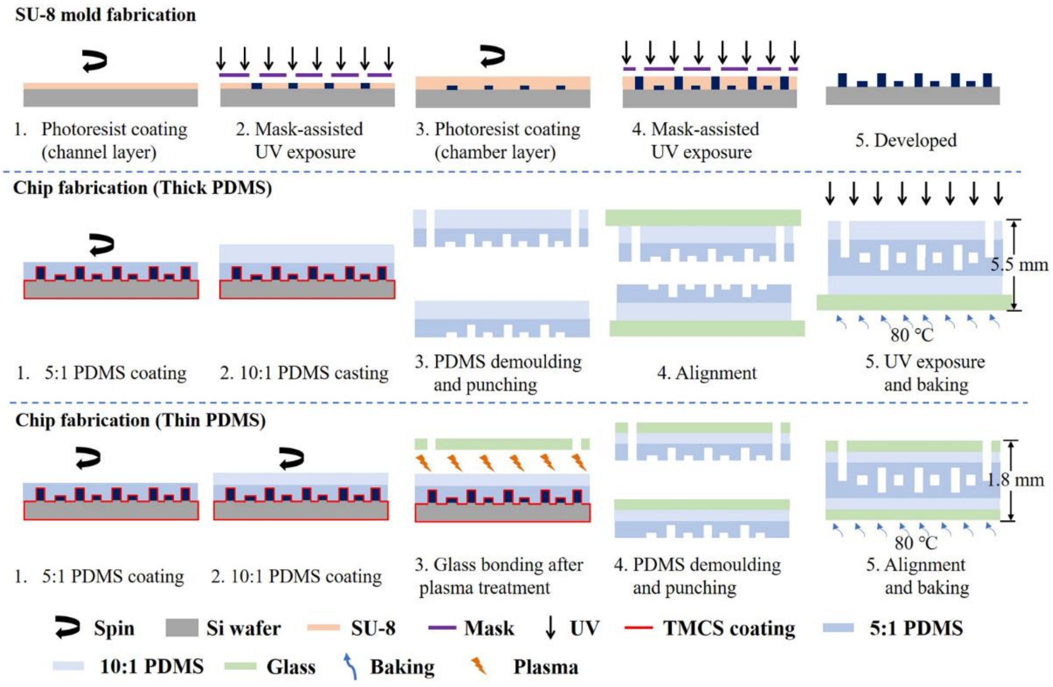


**Figure S18**. Flow chart of chip fabrication.
